# Activation of IL-17+ ILC subsets in IL-18R-deficient mice during fungal allergen exposure

**DOI:** 10.1101/2025.07.17.665289

**Authors:** Allyssa N. Strohm, Blake Civello, Jana H. Badrani, Lee Lacasa, Kellen Cavagnero, Alex Portillo, Michael Amadeo, Luay H. Naji, Anthea Leng, Rachel E. Baum, Xinyu Wang, Heather M. McGee, Yung-An Huang, Taylor A. Doherty

## Abstract

Group 2 innate lymphoid cells (ILC2s) are critical players during type 2 inflammation present in most forms of asthma. ILC2s are tissue-resident cells that produce cytokines IL-5 and IL-13 critical to eosinophilic airway inflammation, mucus production, remodeling, and hyperresponsiveness. Though each ILC subset (ILC1s, ILC2s, ILC3s) is identified by specific transcription factors, cell surface receptors and cytokine profiles, functional plasticity between ILC subtypes occurs in various contexts. IL-18/IL-18R loci SNPs are linked to asthma in multiple genome-wide association studies and IL-18 has been shown to promote plasticity in ILC2s. Despite this, little is known about the *in vivo* role of IL-18/IL-18R on ILC2 responses in the lung. Within hours after mice were exposed to the fungal allergy *Alternaria alternata*, airway levels of IL-18 and IL-18 receptor expression increased on ST2+ ILCs. Single-cell RNA sequencing of lung cells from Alternaria-challenged mice showed that *Il18* was largely expressed by alveolar macrophages, while IL-18R was highly expressed in IL-13+ILC2s. Utilizing IL-18 receptor knock-out mice (IL- 18R-/-), we observed increases in IL-17A production from both ST2+ and ST2-negative ILCs compared to control mice. We further observed an early increase in dual production of IL-5 and IL-17A in ST2+ ILCs followed by enhanced lung eosinophilia in the absence of IL-18R. Together, our findings suggest that IL-18 signaling prevents IL-17A production from ILC2s and subsequent eosinophilia *in vivo*. A further understanding of the regulation of ILC plasticity may lead to novel therapeutic targets in the treatment of ILC-driven asthma.

## Introduction

The majority of asthmatics have type 2 airway inflammation which is characterized by eosinophilia, hyperresponsiveness, and remodeling features such as mucus production (1). The type 2 cytokines IL-4, IL-5, and IL-13 were initially attributed to CD4+ T helper 2 (Th2) cells, and thought to promote various features of type 2 inflammation (2). However, over the last 15 years, group 2 innate lymphoid cells (ILC2s) have emerged as critical producers of IL-4, IL-5, and IL-13 during type 2 immune responses (3–6). ILC2s are members of the broader innate lymphoid cell (ILC) family, which include ILC1s and ILC3s. All ILCs lack antigen receptors that define T and B cells (7). Classically, ILC1s express the transcription factor T-bet and produce IFN-𝛾 and tumor necrosis factor-alpha (TNFα), ILC2s express GATA3 and produce IL-4, IL-5, and IL-13, while ILC3s express ROR𝛾t and produce IL-17A and IL-22 (8). After mice inhale the fungal allergen *Alternaria alternata*, IL-33 release from epithelial cells leads to rapid and robust ILC2 activation and type 2 lung inflammation independent of adaptive immunity (9–11).

Although ILCs are differentiated by their distinct programs, recent studies have shown functional plasticity between mature ILC subsets. One study demonstrated that a hybrid subset of pathogenic ST2+ ILC2s that produced both IL-13 and IL-17A promoted increased lung eosinophilia after exposure to protease allergen (12). These ILCs are referred to as inflammatory ILC2s. Further, IL-12 and IL-18 promote lung ILC2s to develop characteristics of ILC1s through decreased GATA3 expression, increased T-bet expression, and IFN-𝛾 production (13, 14). The IL- 18/IL-18R axis is of particular interest in asthma as genome-wide association and machine learning studies have demonstrated that both IL-18 and IL-18R are linked to asthma, including severe asthma (15, 16). Further, in patients with allergic rhinitis, an IL-18 polymorphism is associated specifically with *Alternaria* allergy (17). Though some studies suggest that IL-18 reduces ILC2 programs (13, 14), other studies support that IL-18 may activate ILC2s including gaining pathogenic ILC3-like functions (18, 19). Importantly, before anti-IL-18 therapeutics are considered for treating severe asthma, further studies are needed to characterize the *in vivo* phenotype and effect of IL-18/IL-18R signaling on ILC2 activation in asthma models.

In addition to ILC2s, IL-18 affects multiple immune and structural cell populations that are active in lung inflammation (20). Originally known as IFN-𝛾-inducing factor, IL-18 is a cytokine produced by macrophages and epithelial cells which binds to the IL-18R𝛼/IL-18R𝛽 complex to activate T helper 1 (Th1) cells to release IFN-𝛾 (21, 22). Aside from type 1 responses, IL-18 acts together with IL-3 to induce production of type 2 cytokines IL-4 and IL-13 by basophils (23). Similarly, co-administration of IL-2 and IL-18 induced type 2 cytokine production from CD4+ T cells (24). In a conventional allergen challenge model with ragweed extract, IL-18 administered at sensitization increased eosinophilic lung inflammation but had the opposite effect if given during late challenges (25). However, the effect of IL-18R signaling in ILC2 responses during mucosal allergen-induced lung inflammation is unknown.

Here, we characterized IL-18/IL-18R signaling in ILCs after administration of the fungal allergen *Alternaria alternata* intranasally (IN) to wild-type (WT) mice, and also compared ILC activation and tissue pathology after administration of *Alternaria alternata* in WT vs. IL-18R-/- mice. We found that IL-18 mRNA and protein was rapidly induced in lung lysates and IL-18R expression was upregulated on ILC subsets after *Alternaria* challenge. IL-18R-/- mice showed increased lung eosinophilia and ILC IL-17 production. Together, this supports a protective role of IL-18 in the accumulation of pathogenic IL-17+ ILCs during innate type 2 lung inflammation.

## Materials and Method

### Mice

Female C57BL/6J mice aged 6-12 weeks old were obtained from Jackson Laboratories (Bar Harbor, ME). WT mice were age and sex-matched to the IL-18R-/- mice. IL-18R-/- mice were obtained from Dr. Hal Hoffman at UC San Diego (originally from Dr. Richard Flavell, Yale) and bred in house with a C57BL/6 background. All studies are approved through the University of California, San Diego Institutional Animal Care and Use Committee.

### Lung Inflammation Models

Mice were intranasally challenged with 50𝜇g *Alternaria alternata* extract (Greer, Lenoir, NC) diluted in PBS and challenged daily over either 1, 2, or 3 days prior to BAL and lung analysis at 1hr, 3hr, 12hr or 24hr after last challenge.

### BAL and Lung Processing

Bronchoalveolar lavage (BAL) fluid was collected using 2% bovine serum albumin (BSA) (Sigma, St Louis, MO). Supernatant from the first BAL was collected and stored at -20℃ for future ELISA analysis. The whole lung was collected into RPMI and promptly dissociated into a single-cell suspension using the Miltenyi Lung Digestion Kit and Gentle MACS Tissue Dissociator per the company’s protocol (Miltenyi Biotec, Bergisch Gladbach, Germany). Cells were counted using the Novocyte flow cytometer based on size and granularity (ACEA, San Diego, CA).

### Flow Cytometry

One million cells from the lung and BAL were stained with antibodies against surface markers. For intracellular staining, 2 million cells from the lung were stained with antibodies against transcription factors and cytokines. Fc receptors were blocked with CD16/32 (Biolegend, San Diego, CA) before antibody staining. Neutrophils were gated as CD45.2+Siglec F-GR1+ and eosinophils were gated as CD45.2+SiglecF+CD11c- cells, with CD45.2 (PerCP), CD11c (FITC), GR-1 (APC), and Siglec-F (PE). ILCs were gated as CD45.2+Lineage-Thy1.2+ lymphocytes and conventional ILCs were gated as CD45.2+Lineage-Thy1.2+ST2+ lymphocytes with CD45.2 (PerCP), Lineage cocktail (FITC), Thy1.2 (Pacific Blue), and ST2 (APC). The lineage cocktail includes CD3, Gr1, CD11b, CD45R, Ter-119, CD11c, NK1.1, CD5, FceR1, TCR𝛽, and TCR𝛾𝛿. For nuclear and intracellular stains, cells were first surface stained as described above, and then permeabilized using the FoxP3 kit (ThermoFisher, Watham, MA) for nuclear stains and stained for Ki-67 (PE). For intracellular stains to assess cytokines, cells were cultured for three hours with cell-stimulation cocktail and a protein transport inhibitor (ThermoFisher, Watham, MA) at 1 million cells per well. Cells were then surface stained and permeabilized using the BD kit (BD Biosciences, La Jolla, CA) and stained for IL-5 (PE), IL-13 (PE), and/or IL-17A (AmCyan). All flow cytometry experiments were performed using the Novocyte 3000 (Agilent, Santa Clara, CA) and data was analyzed using FlowJo software (Tree Star, Ashland, OR) as we have previously reported (26).

### ELISA

BAL samples stored at -20℃ were analyzed using a mouse IL-33 Duoset ELISA kit per R&D protocols (R&D Systems, Minneapolis, MN). Samples stored at -20℃ were analyzed using a mouse IL-18 ELISA kit per Invitrogen’s protocols (Invitrogen, Carlsbad, CA). Plates were read using a microplate reader model 680 (Bio-Rad Laboratories, Hercules, CA). Microsoft Excel and PRISM by GraphPad (San Diego, CA) were used to analyze ELISA results.

### Single-cell RNA sequencing

Female C57BL/6 mice were intranasally challenged with *Alternaria alternata* for 3 days. One day after the last challenge, the mice were euthanized by CO2 inhalation followed by lung isolation. The lungs were collected for single-cell suspension and ILC enrichment (by Mouse Pan-ILC Enrichment Kit; STEMCELL Technologies, Vancouver, British Columbia, Canada). ILCs were enriched by Mouse Pan-ILC Enrichment Kit (STEMCELL Technologies, Vancouver, British Columbia, Canada) and preserved in PIPseq^TM^ T100 3ʹ Single Cell Capture and Lysis Kit prior to processing by sequencing service (Fluent BioSciences, Watertown, MA). The barcode-gene matrices from PIPseeker pipeline were analyzed with the R package Seurat (v4.3.0) (27). Datasets underwent Seurat’s workflow of SCtransform normalization and integration to generate the primary clustering at the resolution of 0.3. Uniform manifold approximation and projection (UMAP) was employed for dimensionality reduction and visualization of the data. For deeper ILC investigation, the clusters with positive expression of *Ptprc* and *Thy1* were identified and with secondary clustering at a resolution of 1.0. Our publicly available single-cell RNA sequencing data (GEO: GSE261844) was analyzed for sources and levels of *Il18*, *Il18rap,* and *Il18r1* (28).

### Statistical Analysis

Statistical Analysis was performed with PRISM Software (Graphpad Software, La Jolla, CA). P-values less than 0.05 was considered statistically significant and were obtained using unpaired t test. *p<0.05, **p<0.01, ***p<0.001.

## Results

### Airway IL-18 and ILC IL-18R are induced by Alternaria exposure in WT mice

To determine the role of IL-18 in a lung allergen model of type 2 inflammation, WT mice were intranasally (IN) challenged with 50𝜇g of *Alternaria.* At 1, 3, and 12 hours (HR) post-challenge, BAL was collected for ELISA to measure IL-33 and IL-18. IL-33 increased rapidly within an hour of IN *Alternaria* challenge as expected and decreased by 3 hours, while being reduced to baseline levels by 12 hours (Figure 1A) (9–11, 29). IL-18 also increased by 1 hour after challenge but remained elevated at 12 hours (Figure 1B). Our recently published single-cell RNA sequencing data of lung cells from *Alternaria*-challenged mice (28) was analyzed for expression of *Il18* in CD45+ leukocytes (Figure 1C) and non-immune cells (CD45-; Figure 1D). Approximately 60% of alveolar macrophages highly expressed *Il18* (Figure 1C). About 5% of fibroblasts and ciliated epithelial cells also expressed *Il18* (Figure 1D).

**Figure 1.**
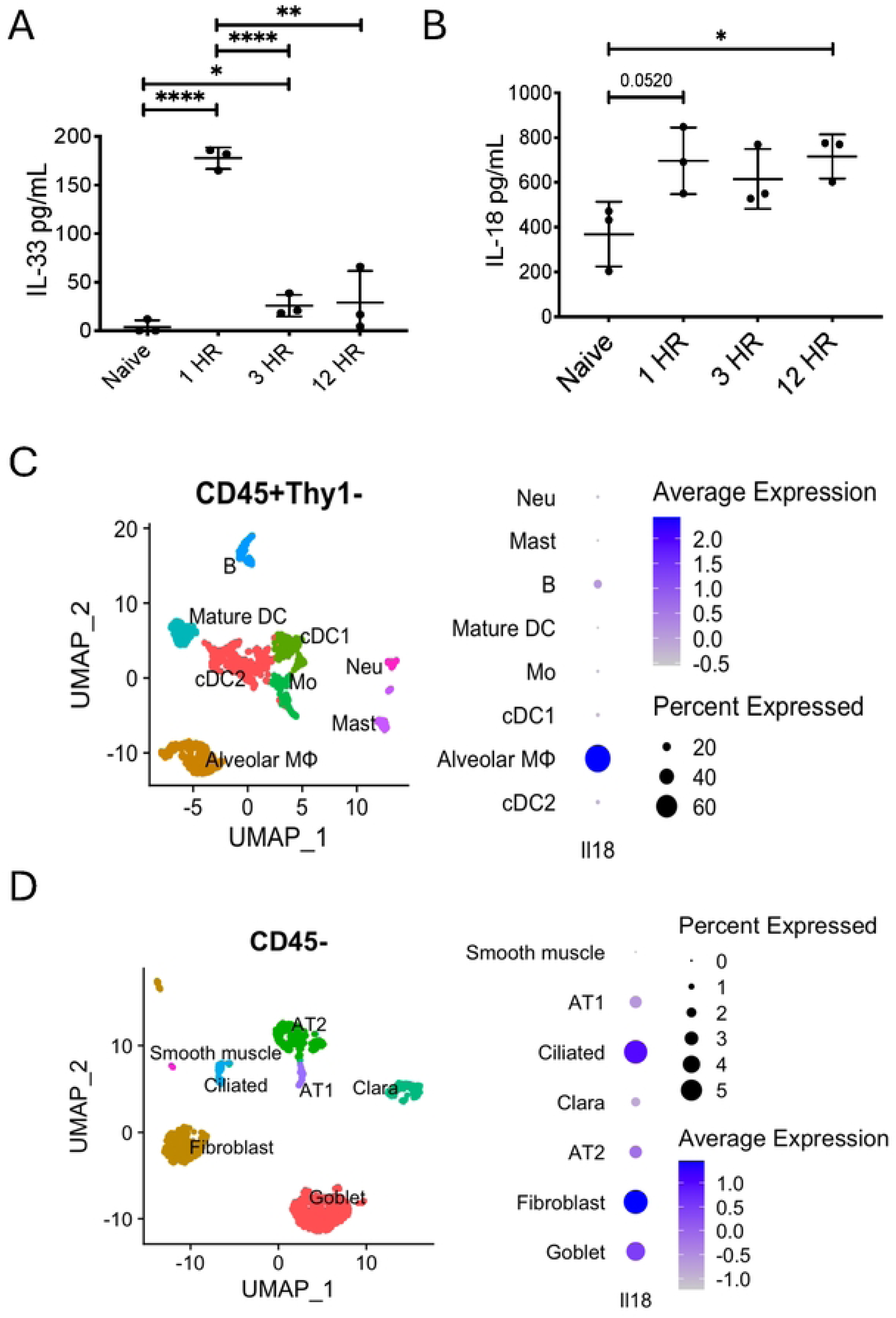
Airway IL-18 is induced by Alternaria exposure in WT mice. WT mice were challenged with *Alternaria alternata.* 1, 3, and 12 hours after intranasal challenge, BAL was collected for ELISA analysis to measure (A) IL-33 and (B) IL-18. Unpaired t test, *p<0.05, **p<0.01, ***p<0.001. *Il18*-expressing cell types identified by scRNA-seq from female WT B6 mice challenged for 3 days with *Alternaria alternata*. The cells that either express (C) *Ptprc* but without *Thy1*:CD45+Thy1- or (D) negative in *Ptprc*: CD45- were exhibited with uniform manifold approximation and projection (UMAP). The expression of *Il18* in (C) CD45+Thy1- or (D) CD45- was presented as dot plot.

ILCs are rapidly activated after airway *Alternaria* exposure in mice and can respond to IL- 18 in various contexts (13, 18, 19, 30, 31). Single-cell RNA sequencing data of enriched lung ILC2s from *Alternaria*-challenged mice (28) showed that IL-18 receptor (IL-18R) subunits IL- 18Rα (encoded by *Il18ra*) and IL-18Rβ (encoded by *Il18rap*) were expressed in CD45+Thy1+ ILCs (Figure 2A). IL-18R transcripts were highly expressed in NK/ILC1s as expected but also significantly upregulated in the IL-13+ ILC2 cluster (Figure 2A). Expression of the IL-18 receptor (IL-18R) at a protein level was assessed in the total Lin-negative Thy1.2+ ILC population by flow cytometry. The percent of IL-18R+ILCs from challenged mice increased along with expression levels of IL-18R (Figure 2B). We next determined the percent of IL-18R-expressing ST2+ILC2s and ST2-negative ILC subsets in naïve and challenged mice. IL-18R+ ST2+ ILC2s increased nearly 5-fold after challenge compared with ST2-negative ILCs that only slightly increased after challenge (Figure 2C). Thus, *Alternaria* challenge in wild type mice leads to rapid increases in airway IL-18 levels and ILC expression of IL-18R, particularly in the ST2+ILC2 population.

**Figure 2.**
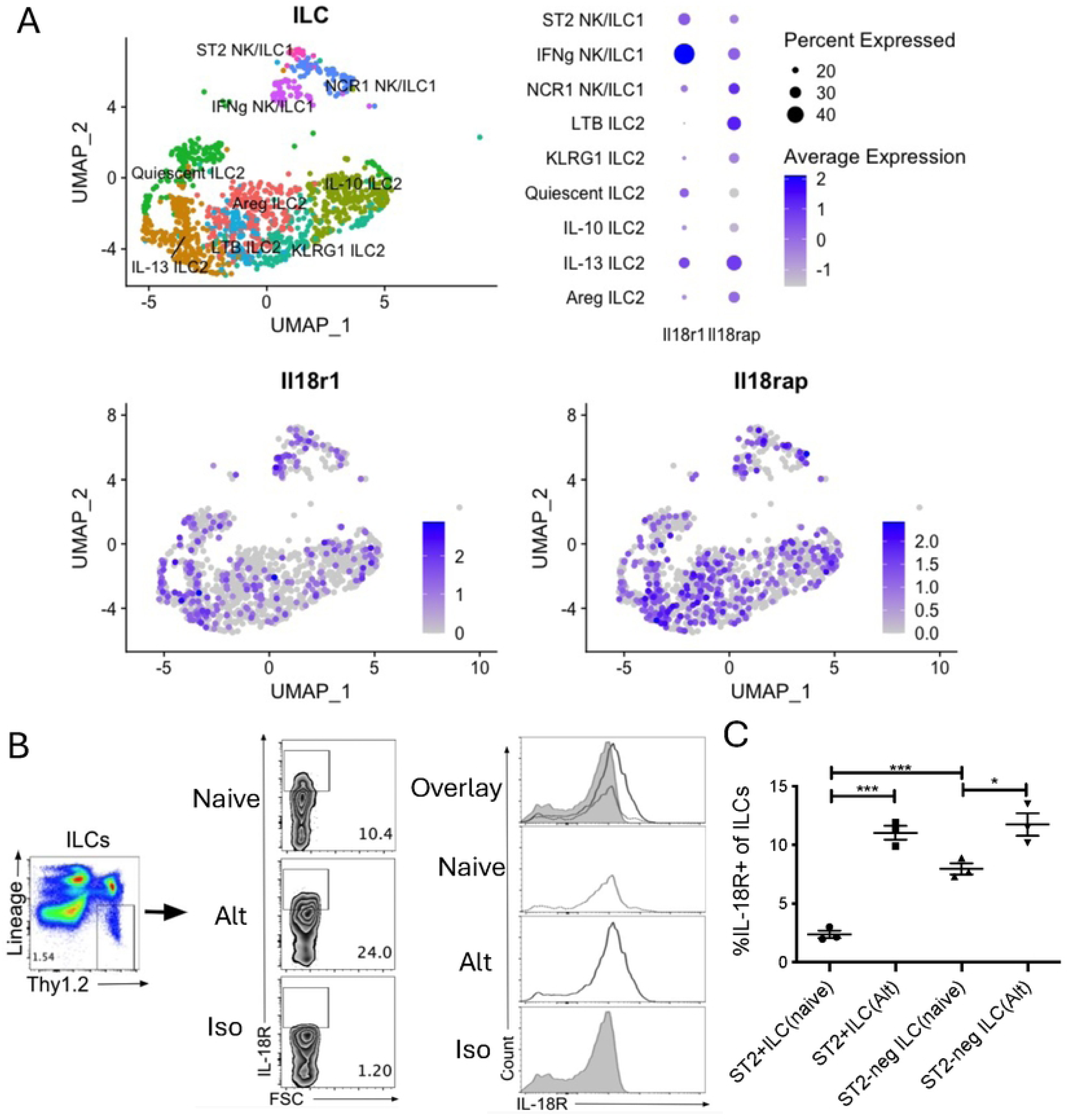
*ILC IL-18R is induced by Alternaria exposure in WT mice*. IL-18R gene expression in (A) *Ptprc*+*Thy1*+ ILCs were identified by scRNA-seq from female WT B6 mice challenged for 3 days with *Alternaria alternata*. Gene expression of IL-18Rα (*Il18r1*) and IL-18Rβ (*Il18rap*) was presented as dot plot (upper) and feature plots (lower). WT mice were challenged with *Alternaria alternata* for three days and lung cells were collected for analysis of ILC IL-18R expression by flow cytometry and compared to unchallenged naïve mice. Representative (B) flow plots and histograms of IL-18R+ lung ILCs. Contains data from two separate experiments. (C) Frequency of IL-18R+ST2+ and IL-18R+ST2- ILCs within the lung of challenged and naïve mice. Unpaired t test, *p<0.05, **p<0.01, ***p<0.001.

### IL-18R signaling suppresses lung eosinophilia after Alternaria challenges

To determine the role of IL-18R in vivo after *Alternaria* challenges, we compared the Siglec-F+CD11c-negative eosinophil levels in lung and BAL of IL-18R-/- mice and WT mice challenged with 50𝜇g of *Alternaria* IN *daily* for three days. Eosinophils in both the BAL (Figure 3A&C) and lung (Figure 3B&D) were elevated on days 1, 2, and 3 in IL-18R-/- mice compared with wild-type controls. In contrast, there was no significant difference in neutrophils in IL-18R-/- compared to WT mice (Supplemental Figure 1). Thus, IL-18R negatively regulates lung eosinophilia throughout the first three days after fungal allergen challenges.

**Figure 3.**
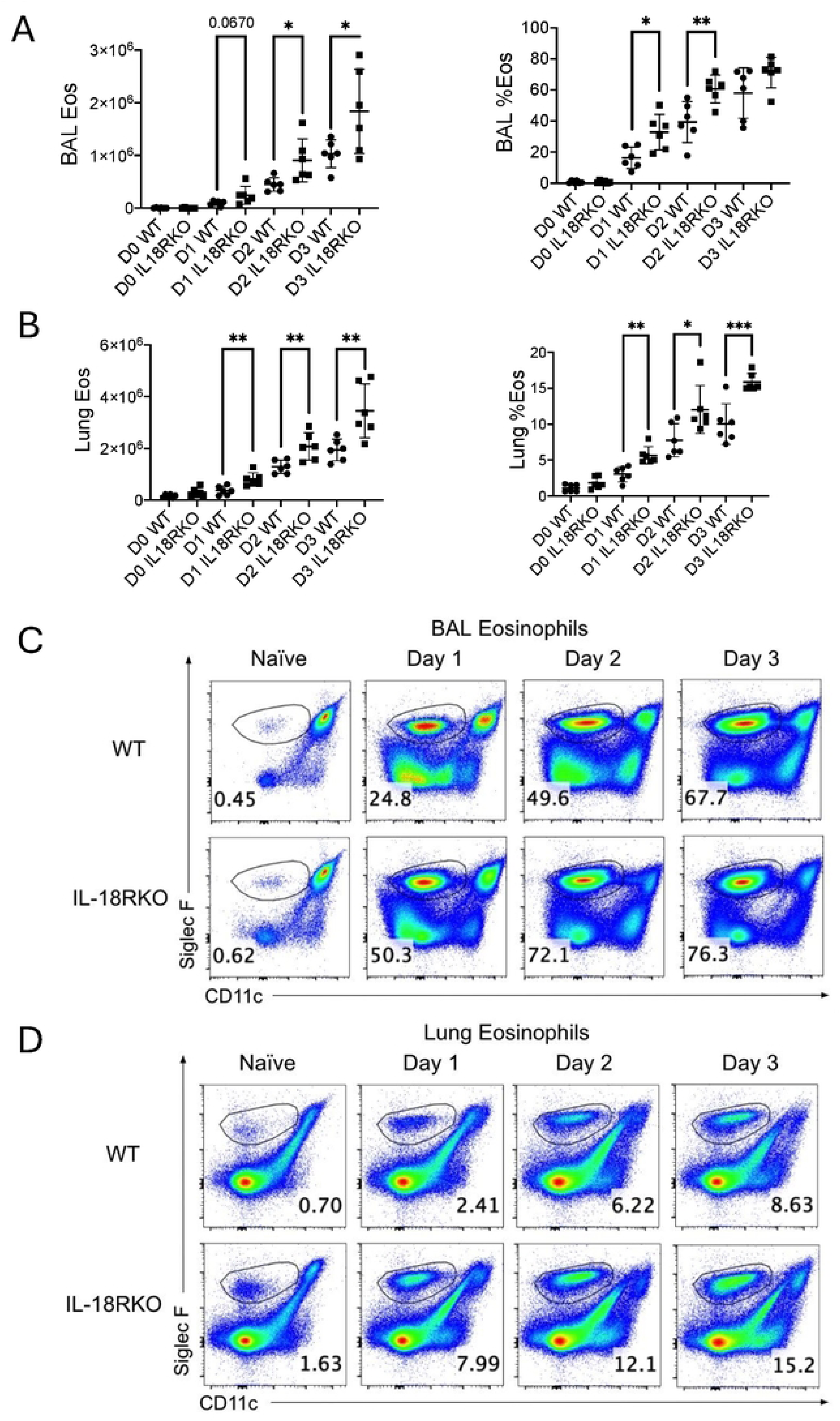
*IL-18R signaling suppresses lung eosinophilia after Alternaria challenges* WT and IL-18R-/- mice were challenged with *Alternaria alternata* for 1, 2 or 3 days, and BAL and lung cells were collected 1 day after the last challenge. (A) Numbers and frequency of Siglec F+CD11C-negative eosinophils in the BAL. (B) Numbers and frequency of eosinophils in the lung. (C) Representative flow plots of eosinophils in the BAL. (D) Representative flow plots of eosinophils in the lung. Unpaired t test, *p<0.05, **p<0.01, ***p<0.001

Since ILC2s are known to recruit eosinophils through type 2 cytokine expression in the innate *Alternaria* model (32), we compared lung ST2+ and ST2-negative ILC frequencies in challenged IL18R-/- and WT mice. There were no differences in the percentages or total numbers of ILC subsets (Figure 4A & B). There was a modest increase in ST2-negative ILC numbers after two days of *Alternaria* challenge, but no change in ST2+ ILCs (Figure 4C & D). Despite the lack of differences in total ILC subset numbers, the percent Ki67+ proliferating IL-18R-/- Lin-Thy1.2+ ILCs were increased on day 1 (Supplemental Figure 2A & B). The number and frequency of Ki67 expression in ST2-negative ILCs from IL-18R-/- mice was increased on days 1 and 2 compared to WT mice, while ST2+ ILCs showed a significant increase in % Ki67 expression on day 1 alone (Supplemental Figure 2C & D).

**Figure 4.**
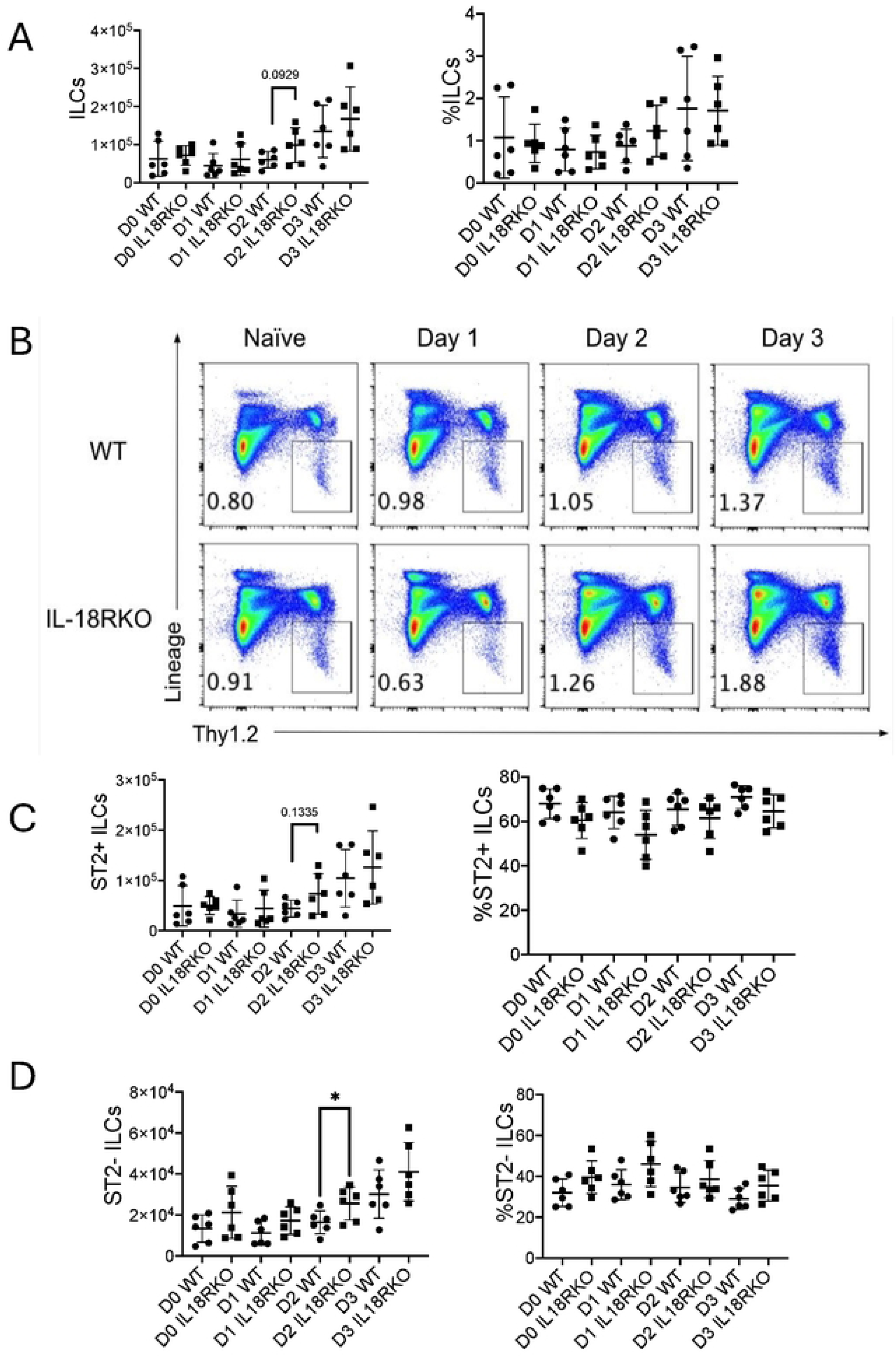
*ST2+ and ST2-negative ILCs unchanged in challenged IL-18R-/- mice* WT and IL-18R-/- mice were challenged with *Alternaria alternata* for either one, two, or three days, and lung cells were collected one day after the last challenge. (A) Number and frequency of total CD45+ Lin-negative Thy1.2+ ILCs in the lung. (B) Representative flow plots of total lung ILCs. (C) Number and frequency of ST2+ ILCs in the lung. (D) Number and frequency of ST2- negative ILCs in the lung. Unpaired t test, *p<0.05, **p<0.01, ***p<0.001

### IL-17A, but not IL-5, expressing ILCs are negatively regulated by IL-18R

Although there were no appreciable differences in the number of ILCs during the 3-day challenge model, we reasoned that differences in ILC cytokine expression may instead be different between IL-18R-/- and WT mice. Surprisingly, despite the observed increased eosinophilia in IL- 18R-/- mice, we did not observe any difference in IL-5+ Lin-Thy1.2+ ILCs between groups of mice at any time point (Figure 5A & B). Moreover, neither the ST2+ nor the ST2- ILC sub- populations frequencies showed changes in IL-5 production (Figure 5C & D). Similarly, we did not identify any differences in IL-13 production between IL-18R-/- and WT mice (Supplemental Figure 3).

**Figure 5.**
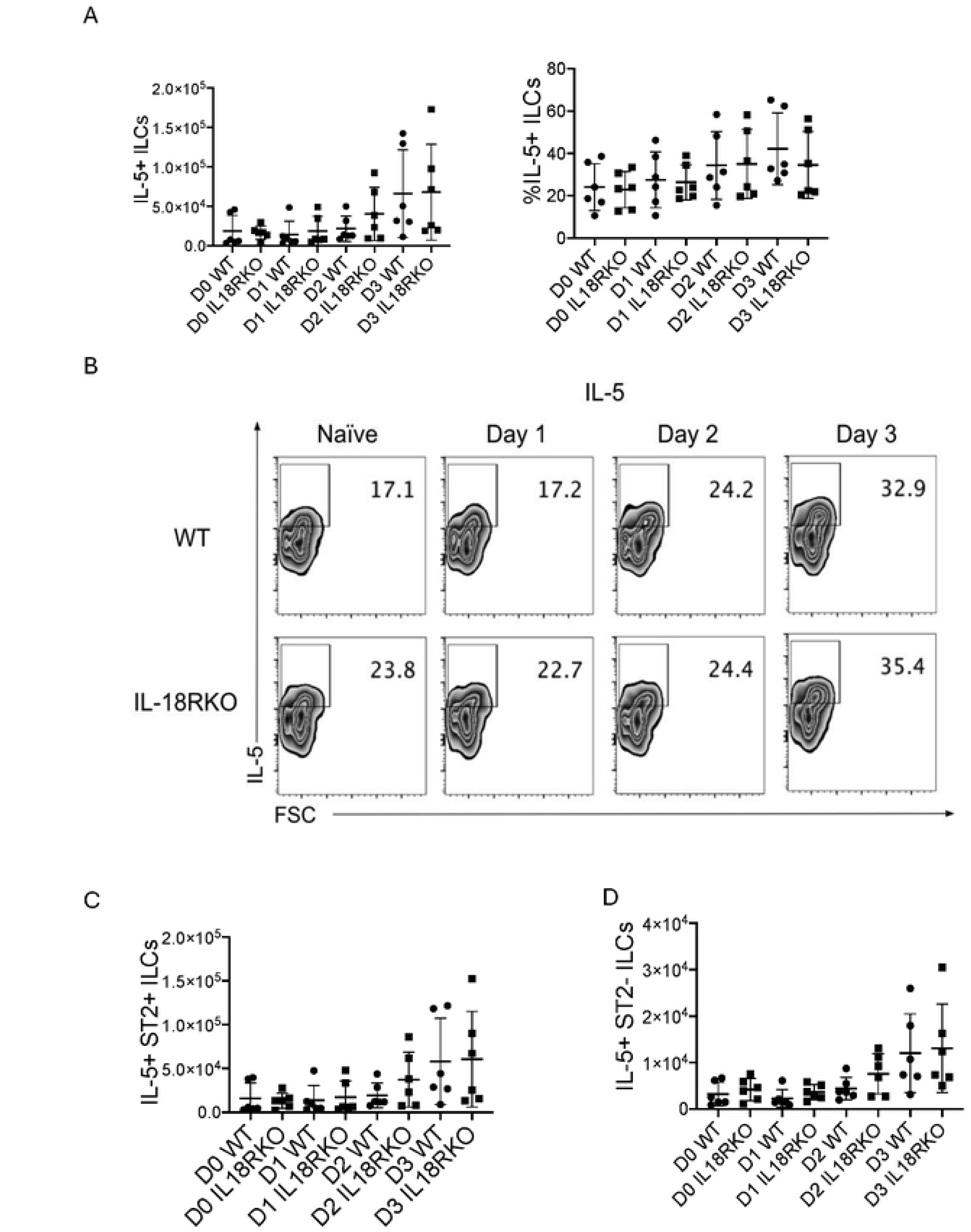
IL-5+ *ST2+ and ST2-negative ILCs unchanged in challenged IL-18R-/- mice* WT and IL-18R-/- mice were challenged with *Alternaria alternata* for either one, two, or three days, and lung cells were collected one day after the last challenge. CD45+ Lin-negative Thy1.2+ ILCs were stained for ST2 and intracellular IL-5. (A) Number and frequency of IL5+ ILCs in the lung. (B) Representative flow plots of total IL-5+ ILCs. Number of (C) IL-5+ST2+IL-5+ and (D) ST2-negative lung ILCs. Unpaired t test, *p<0.05, **p<0.01, ***p<0.001

As inflammatory ILC2 populations expressing IL-17 exist and could exacerbate eosinophilic lung inflammation (12, 19, 26, 33), we assessed levels of L-17A+ ILC subsets. Interestingly, the total numbers of IL-17A+ ILCs were elevated in IL-18R-/- mice at all time points, including at baseline (Figure 6A). The frequency of IL-17A+ ILCs was significantly increased on Day 1 (Figure 6A & B) and showed a trend toward an increase in IL-18R-/- mice compared to WT mice on days 0, 2, and 3. The percentage ST2+ IL-17A+ ILCs from IL-18R-/- mice showed the largest percent increase at all time points compared to control mice, but there was only an increase in IL-17+ ST2+ ILC numbers on Day 0 and Day 2 (Figure 6C, Supplemental Figure 4A). The percent of total IL-17A+ ST2-negative ILCs in IL-18R-/- mice increased on days 1-3, though the percent of IL-17A expression in the ST2-negative population showed only a trend toward an increase (Figure 6D, Supplemental Figure 4B).

**Figure 6.**
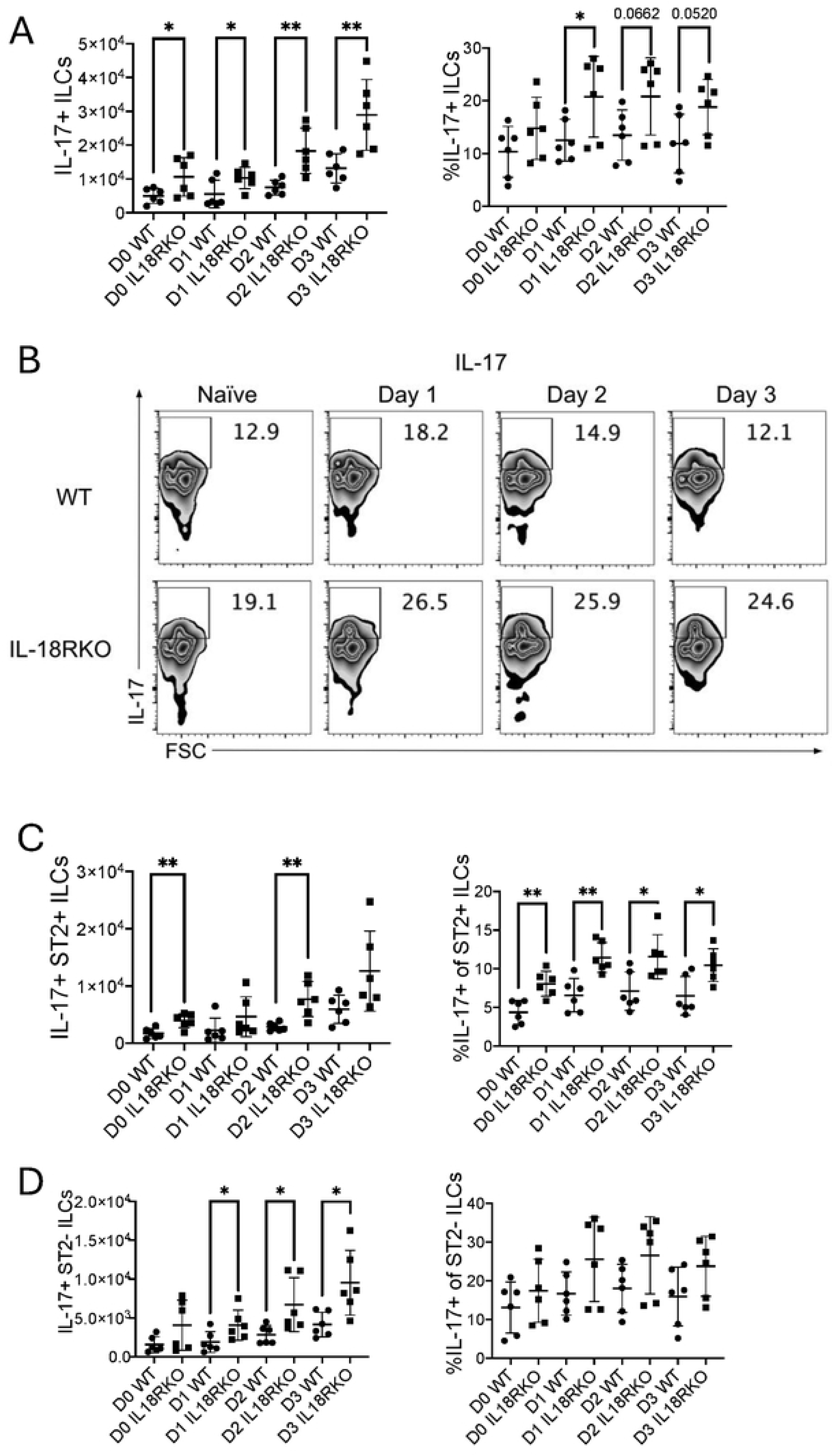
*Increased IL-17A-expressing ILCs in IL-18R-/- mice* WT and IL-18R-/- mice were challenged with *Alternaria alternata* for either one, two, or three days, and lung cells were collected one day after the last challenge. CD45+ Lin-negative Thy1.2+ ILCs were stained for ST2 and intracellular IL-17A. (A) Number and frequency of IL-17A+ total ILCs in the lung. (B) Representative flow plots of IL-17A+ total ILCs in the lung. (C) Number and frequency of IL-17A+ST2+ ILCs in the lung. (D) Number and frequency of IL-17A+ST2-negative ILCs in the lung. Unpaired t-test, *p<0.05, **p<0.01, ***p<0.001

We next assessed dual expression of IL-5 and IL-17A in ILC populations after *Alternaria* challenged IL-18R-/- and WT mice (34). IL-18R-/- mice had significantly higher IL-5 + IL-17A producing lung ILC (IL-5+IL-17A+ ILC) numbers and frequency on day 1 compared to WT mice (Figure 7A & B). Interestingly, ST2+IL5+IL-17A+ ILCs were increased in IL-18R-/- mice at baseline suggesting homeostatic control of this population. The percentage of IL-5+ IL-17A+ ST2+ILC population also increased one day after challenge in IL-18R-/- mice but there was only a trend toward an increase in the absolute numbers of IL-5+IL-17+ ST2+ ILCs, despite there being a statistically significant increase in the number of IL-5+ IL-17+ ST2+ ILCs at baseline (day 0) in IL-18R-/- mice (Figure 7C & D). At day 1 post-challenge, the ST2-negative ILC population showed a significant increase in the frequency and number of total IL-5+IL-17A+ ILCs in the IL- 18R-/- mice (Figure 7E-F). Given that IL-17A is a key cytokine produced by ILC3 and ILC3-like cells, our data suggests an early and possibly homeostatic, role for IL-18R in controlling activated lung ILC2-IL17 / ILC3-like cells.

**Figure 7.**
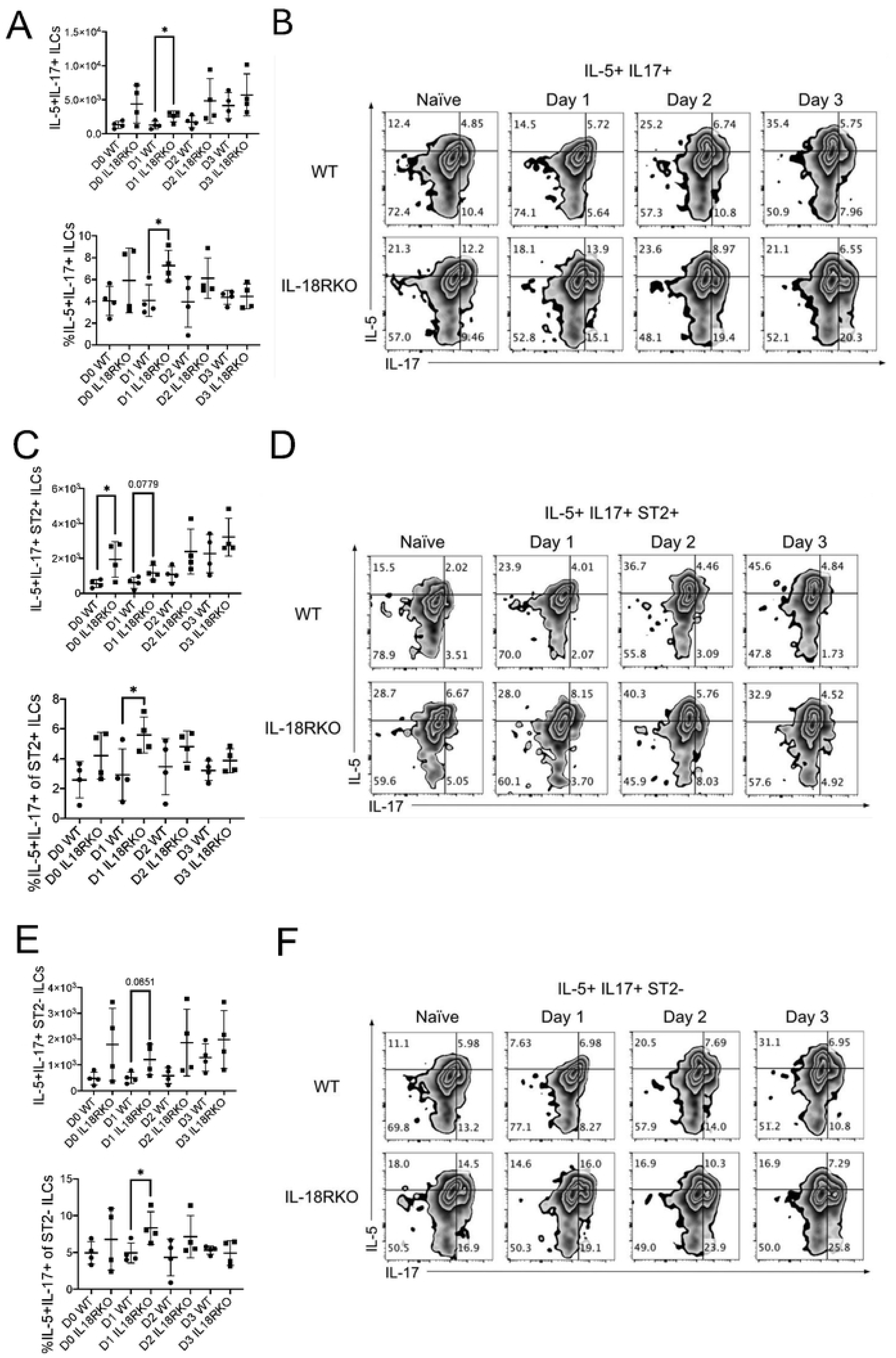
*Early increased dual IL-17A/IL-5-expressing ILCs in IL-18R-/- mice* WT and IL-18R-/- mice were challenged with *Alternaria alternata* for either one, two, or three days, and lung cells were collected one day after the last challenge. CD45+ Lin-negative Thy1.2+ ILCs were stained for ST2 and intracellular IL-17A and IL-5. (A) Number and frequency of IL- 5+IL-17A+ total ILCs in the lung. (B) Representative flow plots of IL-5+IL-17A+ total ILCs in the lung. (C) Number and frequency of IL-5+IL-17A+ ST2+ lung ILCs. (D) Representative flow plots of IL-5+IL-17A+ ST2+ ILCs. (E) Number and frequency of IL-5+IL-17A+ ST2-negative ILCs in the lung. (F) Representative flow plots of IL-5+IL-17A+ ST2-negative ILCs in the lung. Unpaired t-test, *p<0.05, **p<0.01, ***p<0.001

## Discussion

Upon activation, ILC2s critically drive respiratory type 2 inflammatory inflammation through the production of type 2 cytokines (35, 36). While the subclassification of ILCs is based on transcription factor and cytokine production, there is significant ILC heterogeneity as evidenced by “hybrid” ILCs and specific cytokine milieus that can induce ILC plasticity (37–39). In our study, we determined that the fungal allergen *Alternaria alternata*, which is associated with severe asthma (40, 41), rapidly induces airway IL-18 production. Expression of IL-18R was increased on lung ILCs after *Alternaria* exposure and IL-18R-/- mice showed early enhanced eosinophilic lung inflammation. Surprisingly, ILC production of IL-17A *in vivo* was primarily negatively regulated by IL-18R.

IL-18 and IL-18R have long been linked to asthma including severe asthma (16, 42–47). Previous studies demonstrated that *Alternaria* induces human and mouse airway epithelial IL-18 production (48, 49). Our findings are consistent with reports demonstrating increases in lung IL- 18 after *Alternaria* exposure and support a model by which IL-18 may contribute to fungal asthma. Our scRNA-seq data suggests that the dominant source of IL-18 transcripts is alveolar macrophages compared with epithelial cells or other non-immune cells. Previous studies have shown that macrophages are a known source of IL-18 which is processed by NLRP3 and contributes to acute lung inflammation in multiple models including cigarette smoke-induced lung disease (50, 51).

Our group and others have previously shown that *Alternaria* rapidly activates lung ILC2s (10, 11, 52, 53). Previous studies by other groups have investigated the *in vitro* effects of IL-18 on ILC2s, showing an increased activation or induction of plasticity (13, 18, 19, 30, 31). However, the effects of IL-18 on ILC2 *in vivo* had not been previously elucidated. In the current study, we found that IL-18R mRNA and protein expression were increased on ILC2s after *Alternaria,* suggesting that ILC2s are primed to respond to IL-18. A previous study demonstrated that house dust mite (HDM) allergen increased IL-18R in human CD4+ Th2 and Th17 cells as well as mouse Th2 cells during OVA-induced lung inflammation (54). HDM along with IL-18 was also shown to increase IL-18R in human ILC2s from allergic rhinitis patients (30). However, to our knowledge, no reports have demonstrated that *Alternaria* induces ILC2 expression of IL-18R that could, in addition to Th2 and Th17 cell expression, contribute to broad innate and adaptive IL-18- mediated effects during fungal allergen exposure.

Given the potential role of IL-18 in asthma, especially severe asthma, the possibility of targeting IL-18 has been proposed as a therapeutic (16, 43). As IL-18 has complex effects on multiple cell types (20, 55), we studied the *in vivo* role of IL-18R signaling during type 2 lung inflammation Interestingly, our study demonstrates IL-18R negatively regulated *Alternaria-* induced airway eosinophilia without a detectable suppression of IL-5 production from ILCs. While we did not observe any changes in IL-5 production from IL-18R-/- ST2+ ILCs, there was an increase in IL-17A production from IL-18R-/- ST2+ ILCs. Conventional ST2+ ILC2s don’t typically produce IL-17A, though we found significant increases in IL-5+IL-17A+ co-production from ST2+ ILCs in IL-18R-/- mice. A similar population of “pathogenic/inflammatory” lung IL- 17A+ST2+ ILC2s has been reported during IL-33 and papain exposures (12). Our study suggests that IL-18 specifically regulates the IL-17A production in this population of ILC2s.

Elevated airway IL-17A levels are linked to neutrophilia in asthma, though the precise role of IL-17A in asthma or how IL-17A is regulated by IL-18 are not well characterized (56–59). In a study that examined the effects of IL-18 in intestinal inflammatory diseases, CD4+ T cells that express the IL-18R were found to limit Th17 differentiation, thereby maintaining a homeostatic condition (60). Alternatively, IL-18 binding protein (IL-18BP), used as a decoy receptor for IL- 18, attenuated levels of IL-17A production in a rheumatoid arthritis model (61, 62). Previous reports demonstrated that IL-18 promoted IL-17A secretion in a cecal slurry-induced peritonitis model (63) and, along with IL-23, induced T cells to produce IL-17A in an experimental autoimmune encephalomyelitis model (64). A very recent report demonstrated that IL-18 in combination with IL-1β induced IL-17A production from purified human ILC2s (19). In contrast, we observed an increase in IL-17A production from ST2+ ILCs in *Alternaria*-exposed IL-18R-/- mice, and this change in IL-17A was not associated with a significant change in the type 2 cytokines IL-5 and IL-13. Considering the pathogenic role that IL-17A-producing ST2+ cells may play in refractory asthma (12, 19), future studies will need to better clarify the role of IL-18 in the regulation of this IL-17A+ ST2+ ILC2 population. The results from our study in mice suggest a protective role for IL-18 in the prevention of ILC2s becoming ILC3-like by preventing the production of IL-17A from ST2+ ILCs, which could be context (*in vitro* versus *in vivo*) and species-dependent.

We found an increase in lung eosinophils during *Alternaria* challenge in IL18R-/- mice compared with controls. Several studies have examined the role of IL-18 in the development of eosinophilia and have found that IL-18 plays a direct role in the maturation and induction of eosinophils (65, 66). Thus, we cannot rule out a direct effect of IL-18/ IL-18R signaling on lung eosinophils . However, we did find a significant increase in IL-5+ IL-17A+ ILCs as well as the IL-17A in both the ST2+ and ST2-negative ILC subsets. Interestingly, a previous report demonstrated an important role of IL-17A+ ST2+ ILCs in IL-33-induced lung inflammation and found that blockade of IL-17A reduced airway eosinophilia (12). The mechanism could include the ability of IL-17A to induce the expression eotaxin-1 (CCL11) among other possibilities (67–69). Additionally, Margaret *et al*. demonstrated that a low level of IL-17A in the presence of IL- 13 stimulation led to enhanced eosinophilia which could be relevant in our model (67). Thus, it is plausible that the increased ILC IL-17A production in the IL-18R-/- mice could drive increased lung eosinophilia.

Potential signaling mechanisms by which IL-18 regulates IL-17A, include TRAF6 interaction with STAT3, mediating its ubiquitination (70, 71). Since STAT3 is one of the crucial transcription factors for IL-17A, deactivation of STAT3 may attenuate the production of IL-17A and potentially serve as a protective mechanism (72, 73). Interactions between IL-18 and STAT3 have been shown in a spontaneous arthritis model, where the activation of STAT3 mediated IL- 23-induced IL-17A production from CD4+ T cells (74). Furthermore, when STAT3 was deleted from T cells, their IL-17A production was diminished (75). These findings suggest that IL-18 signaling may prevent IL-17A production via regulation of STAT3 in ILCs. However, we must keep in mind that IL-18 has also been shown to directly interact with the JAK/STAT pathway through STAT3 activation, making it difficult to determine if STAT3 plays a role in IL-18 signaling vs. IL-17A signaling (76).

Overall, our studies support a protective role of IL-18 in an innate model of fungal allergen-induced lung inflammation. We showed that IL-18 is induced by *Alternaria* exposure and attenuates eosinophilia and production of IL-17A production in lung ILCs. While the mechanisms behind the regulatory roles of IL-18 in asthma continue to be characterized, further investigations into the regulation of ILC plasticity may provide insight into asthma pathogenesis and the implications of targeting IL-18R to treat this disease.

